# Identification of new ciliary signaling pathways in the brain and insights into neurological disorders

**DOI:** 10.1101/2023.12.20.572700

**Authors:** Abdelhalim Loukil, Emma Ebright, Akiyoshi Uezu, Yudong Gao, Scott H Soderling, Sarah C. Goetz

## Abstract

Primary cilia are conserved sensory hubs essential for signaling transduction and embryonic development. Ciliary dysfunction causes a variety of developmental syndromes with neurological features and cognitive impairment, whose basis mostly remains unknown. Despite connections to neural function, the primary cilium remains an overlooked organelle in the brain. Most neurons have a primary cilium; however, it is still unclear how this organelle modulates brain architecture and function, given the lack of any systemic dissection of neuronal ciliary signaling. Here, we present the first in vivo glance at the molecular composition of cilia in the mouse brain. We have adapted *in vivo* BioID (iBioID), targeting the biotin ligase BioID2 to primary cilia in neurons. We identified tissue-specific signaling networks enriched in neuronal cilia, including Eph/Ephrin and GABA receptor signaling pathways. Our iBioID ciliary network presents a wealth of neural ciliary hits that provides new insights into neurological disorders. Our findings are a promising first step in defining the fundamentals of ciliary signaling and their roles in shaping neural circuits and behavior. This work can be extended to pathological conditions of the brain, aiming to identify the molecular pathways disrupted in the brain cilium. Hence, finding novel therapeutic strategies will help uncover and leverage the therapeutic potential of the neuronal cilium.

## Introduction

The primary cilium is a sensory organelle that is essential for embryonic development and the homeostasis of multiple tissues and organs. It protrudes from the cell to detect extracellular signals and generates specific downstream responses ^1^. Cilia therefore play critical roles in transducing major signaling pathways, including the Sonic Hedgehog (SHH) pathway ^2–7,1^. Mutations in ciliary genes are associated with multiple recessive genetic conditions, collectively termed "ciliopathies," which are often characterized by severe neurological deficits ^1,8–11^, including cognitive impairment, motor dysfunction, and brain malformations ^12–15^. Recent evidence has linked cilia dysfunction to additional neurodevelopmental and neurodegenerative disorders ^16–23^. While cilia are intimately linked to brain disorders, the mechanisms underlying cilia-driven pathogenicity in neurological disorders are poorly understood. The ciliary signaling pathways underlying these diseases remain unclear.

Robust evidence highlights the requirements for cilia in preserving the structure and survival of neurons *in vitro* and *in vivo* in the adult brain ^24–29^. In a previous study, our laboratory found that cilia ablation in the adult cerebellum caused the degeneration of Purkinje neurons and defective neuronal connectivity ^24^. Other recent work has uncovered structural diversity among cilia of different cell-types in the brain. For example, while neuronal cilia dock directly to the plasma membrane, astrocytic cilia are present in pockets or within the soma ^30,31^. More importantly, the neuronal cilium appears to form a novel tetrapartite structure with pre- and post-synapses and astrocytes in which the cilium has access to synaptic activity in the surrounding environment.^30^. Consistent with these findings, axons were found to release serotonin directly onto neuronal cilia, triggering changes in nuclear actin and increased chromatin accessibility ^29^. Despite such striking findings, the molecular networks within the neuronal cilium that mediate adult brain function and homeostasis are poorly defined. What signals do neuronal cilia detect and process? What are the downstream regulators and their effects on the neuron’s function and structure? In the absence of such knowledge, identifying neuronal circuits that rely on neuronal cilia will likely remain difficult.

The cilium’s sensory function is tightly linked to its protein composition, modulated by a precisely regulated trafficking of molecules in and out of the cilium by intraflagellar transport (IFT) and other associated ciliary membrane trafficking pathways ^32–34^ . Cilia use their intrinsic molecular composition to drive downstream signals and mediate the appropriate cellular response in a tissue-specific manner. The neuronal cilium harbors a number of specialized signaling molecules specifically enriched in neural tissues, including somatostatin receptor 3 (SSTR3), serotonin receptor 5-HT6, and adenylyl cyclase 3 (AC3) ^35–42^ . However, the identification of the cilia proteome in neurons and how it may modulate signaling networks and neural circuitry is still lacking. To date, mammalian ciliary proteomes have been defined mainly through the use of immortalized cell lines, and thus do not account for tissue specificity or the *in vivo* microenvironment. This is particularly true for neuronal cilia, which are present within a neural tissue with complex organization and a variety of cellular subtypes. Recent transmission electron microscopy studies underscore the involvement of neuronal cilia in neural circuitry ^30,31^, which can’t be recapitulated easily in vitro.

In order to address this gap in knowledge, we applied an *in vivo* proximity-dependent biotin identification (iBioID) strategy ^43,44^ to capture a first glance of the proteome of neuronal cilia in the mouse brain, optimizing the experimental conditions that allowed the efficient purification of ciliary proteins from brain tissue. Quantitative mass spectrometry uncovered 389 proteins potentially enriched in neuronal cilia, including 42 known ciliary proteins. Gene ontology analysis revealed that more than 199 genes were associated with or mutated in rare neurological disorders. Using this approach, we have identified novel ciliary signaling nodes in the brain that were not revealed in prior studies of the ciliary proteomes of immortalized cell lines. Based on these findings, we conclude that cilia in neurons are highly enriched with neuropeptide and neuroendocrine receptors, ion channels, and calcium-binding proteins. We also identified several molecular modules involved in cell-cell communication, including axon guidance, the Eph/ephrin pathway, and GABAergic signaling. This work presents a powerful new tool for identifying the *in vivo* proteomes of primary cilia in neurons in the brain.

## Results

### Targeting biotin ligase to neuronal cilia in vivo

To understand how the neuronal cilium mediates neural functions, we focused on defining its intrinsic protein composition in the mouse brain **(Figure 1A, B)**. We deployed *in vivo* BioID^44^, using a ciliary bait to target biotin ligase (BirA or BirA2 ^43,45^) to neuronal cilia without affecting cilia biogenesis and length. We first tested several cilia-targeting baits: the N terminus of NPHP3 (a.a. 1-203) ^46^, the ciliary targeting sequence (CTS) of Fibrocystin ^47^, and the ciliary GPCRs 5-HT6, MCHR1, and SSTR3) ^35–38,40,41^, to which we fused BirA or BirA2 ^43,45^. We initially expressed each of the baits in IMCD3 cells. In the presence of biotin, cells were serum starved for 48 hours to induce cilia formation. We then assessed the subcellular localization of the baits and biotin labeling, as well as their effects on cilia assembly and length **(Figure S1A)**. We next expressed the cilia-BioID constructs in the mouse brain, injecting purified adeno-associated viruses (AAV) coding for either HA-tagged BirA2 (or BirA) alone or each of the cilia-targeting BirA2 (-BirA) into lateral ventricles of neonatal mice at postnatal day (P0) **(Figure 1A**, **S1),** expressed under the *hSynapsin* promoter to drive the transgene’s expression in neurons ^44^. Injected mice recovered for at least 21 days and were then injected with 6.66 M biotin daily for 10 days. At P31-P34, we extracted the mouse brains and generated sagittal sections of the mouse cerebral cortex.

**Figure 1:**
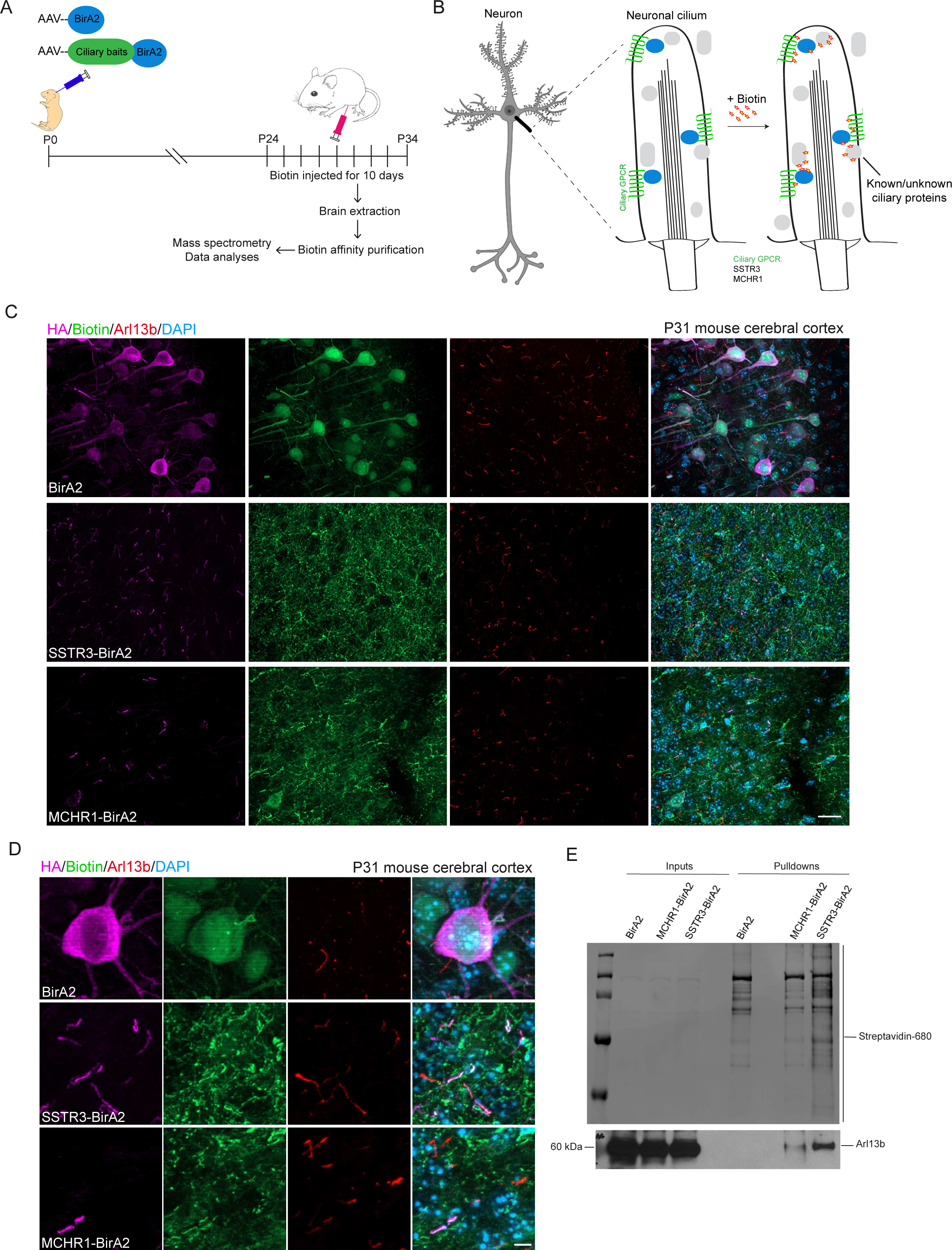
Targeting of the biotin ligase BirA2 to neuronal cilia *in vivo*. **A.** Experimental procedure to label the protein composition of the neuronal cilia in the brain. **B.** A simplified schematic of the targeting strategy for BirA2 using ciliary GPCRs. **C.** Immunohistochemical labeling with antibodies to HA (fused to BirA2, magenta), Biotin (green), and Arl13b (red) on mouse cerebral sections transduced with AAV-BirA2 or AAV-GPCRs-BirA2. DNA was stained with DAPI. Scale bars: 20 µm. **D.** Same as C with zoomed-in images. Scale bars: 5 µm. **E.** Streptavidin pulldowns were performed on total cell lysates from brains transduced with AAV-BirA2 or AAV-GPCRs-BirA2. Western blots were probed with Strepatavidin-680 and Arl13b antibodies.

Sections were labeled with antibodies to biotin, ARL13B, and HA as a readout of the construct’s expression. The most consistent and best performing baits *in vitro* and *in vivo* were the G protein-coupled receptors (GPCRs) MCHR1 and SSTR3, which showed ciliary localization with positive biotin labeling **(Figure 1C, D, and S1)**. Note that BirA2 was fused to the C-terminus of SSTR3 and MCHR1 to localize within the intraciliary lumen. We therefore chose to utilize SSTR3 and MCHR1 as baits, with a focus on mapping the molecular networks within the neuronal cilium. Compared to the MCHR1 bait, SSTR3 had better coverage of the number of cilia labeled, consistent with an overall higher biotinylation profile **(Figure 1C, D, and E)**. Under the control of the hSynapsin promotor ^44^, expression of BirA2 fusions was exclusively found in cells positive for NeuN, a marker for neurons **(Figure S2)**. We also assessed how widespread the biotin labeling was throughout the whole brain for BirA2 alone and GPCRs-BirA2. We found their expression patterns to consistently colocalize with the biotin labeling. The most prominent sections expressing these constructs were from the cerebral cortex **(Figure 1C, D)** and, to a lesser extent, the hippocampus, with limited expression in the cerebellum. We therefore kept the anterior mouse brain and removed the cerebellum for the following experimental procedures.

**Figure 2:**
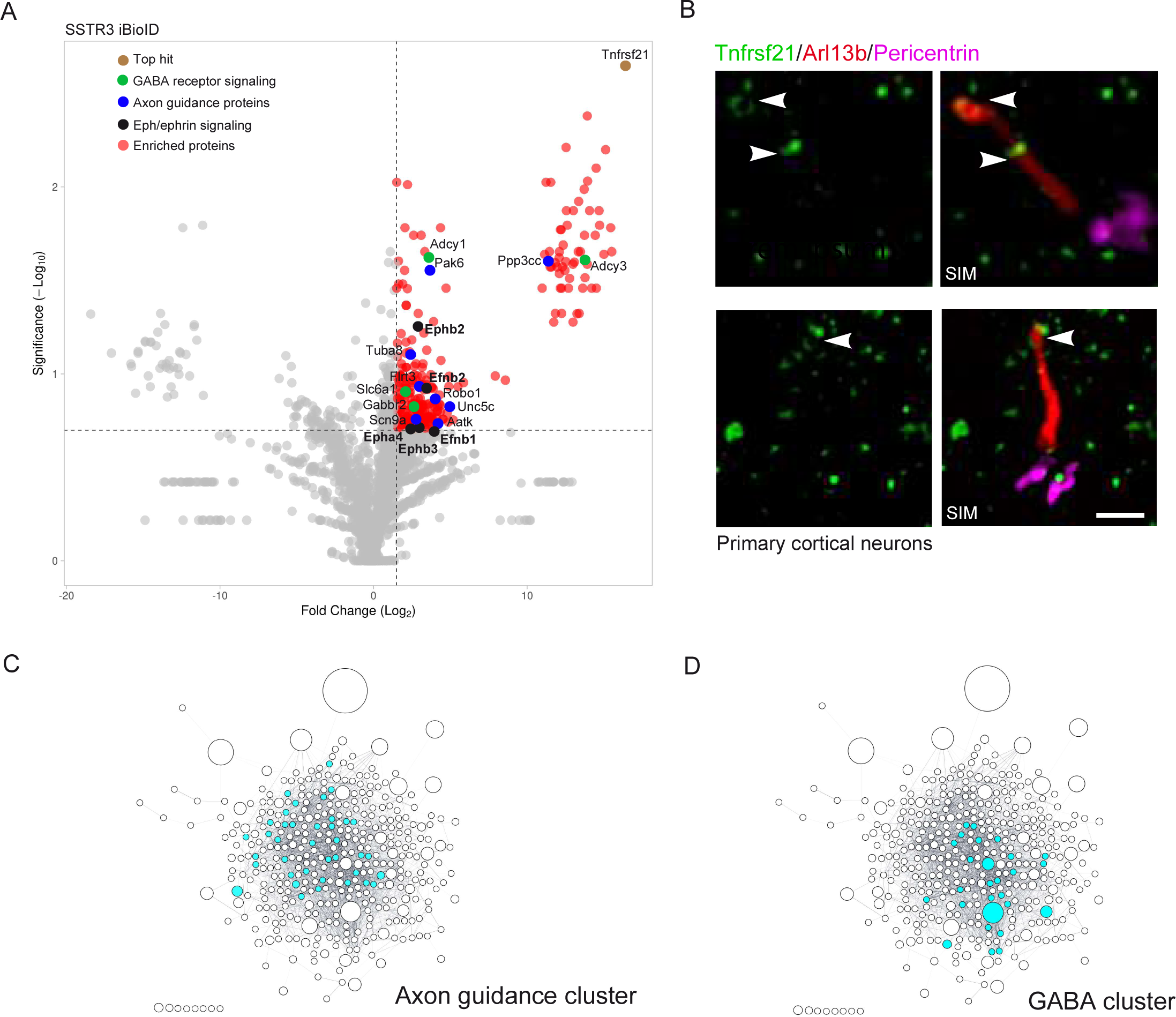
Cilia iBioID identified neural-specific clusters in neuronal cilia *in vivo*. **A.** Volcano plot of SSTR3-proximate proteins labeled by BioID. The red dots show the proteins considered hits, delimited by the selected thresholds: log2 fold change 2 and significance 0.7. The top hit, Tnfrsf21, is highlighted in brown; GABA receptor signaling in green, axon guidance proteins in blue, and Eph/Ephrin signaling in black. The X-axis denotes the log2 fold change of SSTR3-BirA2: BirA2 control. In the Y-axis, significance displays the negative log10 transformed p-value for each protein. **B.** SIM acquisitions show that the top hit, Tnfrsf21, is enriched in the ciliary tip of primary mouse cortical neurons. Two representative examples are shown with cilia by Arl13b (red), centrosomes using Pericenrin (magenta), and Tnfrsf21 (green). Scale bars: 500 nm. **C.** Clustergram topologies of axon guidance hits **D.** GABAergic cluster enriched in Cilia iBioID.

iBioID has been applied primarily to the synaptic compartment of the brain ^44^. To better capture ciliary baits, we partially modified the original iBioID experimental procedure **(Detailed in Materials and Methods)**. Three key changes were made to the prior iBioID: (i) a greater number of AAV particles were injected; (ii) dual-site introduction of biotin solution was combined with longer incubations; and (iii) a less harsh lysis buffer was used during the purification stage. These changes have been key to successfully enriching biotinylated proteins from MCHR1- and SSTR3-BirA2 pulldowns. GPCR-BirA2 pulldowns show a clear enrichment of the ciliary marker ARL13B not observed in the BirA2 alone condition **(Figure 1E)**. Hence, we have found that GPCRs are the most effective baits for our subsequent experiments. The low abundance of ciliary proteins in the brain challenges our ability to enrich ciliary proteins. Our experimental approach addresses this limitation and can be used for low-abundance neural proteomes or cilia subtypes in the anterior mouse brain.

### Neuronal cilia BioID2 unveils novel ciliary nodes with a subset of known ciliary proteins

To obtain a robust set of hits, we performed quantitative mass spectrometry on three biological replicates. We considered as hits all proteins with at least a 2-fold enrichment in GPCRs-BirA2 compared to BirA2 alone. We coupled that with a false discovery rate of at least 0.7 (a p-value of 0.036 or lower) **(Figure 2A)**. A total of 389 proteins were found to be enriched in the SSTR3-BirA2 dataset compared to BirA2 alone **(Figure 2A**, **Table 1)**. Only ten hits were found to be enriched in the MCHR1 dataset **(Figure S3A**, **Table 1)**. This is consistent with insufficiently robust MCHR1 labeling **(Figure 1E)**. Additionally, we identified eight out of ten hits from the MCHR1 iBioID that were found in the SSTR3 iBioID. Hence, we focused our analyses on the SSTR3 iBioID dataset **(cilia iBioID dataset)**. First, we compared our hits with known centrosome-cilia proteomes obtained *in vitro* in mammalian proliferating cells ^46,48^. We uncovered a total of 42 proteins (10.8%), overlapping with our cilia iBioID dataset **(Figure S3B, C, and Table 1)**. These results also suggest overlapping protein repertoires between neuronal cilia and those of proliferating cells. While we used SSTR3, a membrane-associated protein, as bait, we still detected ciliary proteins from other ciliary compartments. This includes early ciliary vesicles as well as the microtubule-base axoneme. Early cilia initiation regulators such as SNAP29, EHD1, EHD3, and RAB8A were enriched in the cilia dataset ^49,50^. Potential hits related to the axonemal compartment have been identified, such as the subunit of kinesin II, KIFAP3, and specific tubulin subunits, including TUBB2A, TUBB2B, and TUBB4B ^51–53^ **(Table 1)**. Modulators of the cilium’s length and stability, such as SEPT6, SEPT7, and SEPT11, were found in our cilia hits ^54^. GPCR downstream effectors such as ADCY3 and ADCY9 were also detected **(Figure S3B, C, and Table1)**. Note that ADCY3 is a well-established marker for neuronal cilia in the brain ^39–42^. We also identified the protein kinase cAMP-activated catalytic subunit beta, PRKACB, which is one of the subunits of the holoenzyme PKA required for Sonic Hedgehog signaling ^55^. Similarly, the non-canonical Wnt target genes PLCB1 and DAAM1 were hits in our cilia dataset ^56^ **(Table 1)**. Overall, our findings from the SSTR3 iBioID covered compartments and signaling pathways previously linked to primary function. This indicates that our updated iBioID experimental procedure successfully captured known ciliary proteins *in vivo* from the brain.

89.2% of the enriched hits were not previously linked to primary cilia. We concentrated on identifying protein networks that could provide insights into novel ciliary nodes driving signaling in the brain. We began validating these findings by examining the subcellular localization of the top hit of cilia iBioID. Tnfrsf21, or death receptor 6 (DR6), is a member of the tumor necrosis factor receptor superfamily and known to mediate axon pruning and neuronal death ^57^. We used primary antibodies against endogenous Tnfrsf21 in mouse primary cortical neurons, co-staining with ARL13B and Pericentrin to label primary cilia and the centrosome, respectively. Using super-resolution microscopy, we have consistently observed Tnfrsf21 in an atypical ring-shaped pattern at the distal cilium (: 63.60% ± 5.97 of cilia positive for Tnfrsf21) **(Fig 2B)**. Such discrete subcellular localization would have been difficult to uncover without our iBioID findings.

### Eph/Ephrin and GABA receptor signaling components are widely enriched in neuronal primary cilia

To identify and prioritize signaling networks enriched in neuronal cilia, we performed an extensive gene ontology (GO) analysis. We first used Gonet ^58^ to determine categories of molecular functions of all cilia hits through GO term enrichment analysis. Each of the categories was generated and linked to the representative genes with a q-value threshold 0.05. **(Table 2)**. Most of the proteins identified are transmembrane transporters, ion channels, and receptors, consistent with SSTR3-BirA2 labeling capturing components in proximity to the ciliary membrane. Using the Ingenuity Pathway Analysis (IPA) database, we found significant enrichment of pathways implicated in cell-to-cell interaction and cellular morphology **(Table 3)**. We also employed Cytoscape to reconstruct the molecular interactome of the cilia dataset. The neuronal ciliary network is constituted by interconnected spheres, in which the size of each sphere is correlated to the fold change value and the color shade to the p-value **(Figure 3)**.

**Figure 3:**
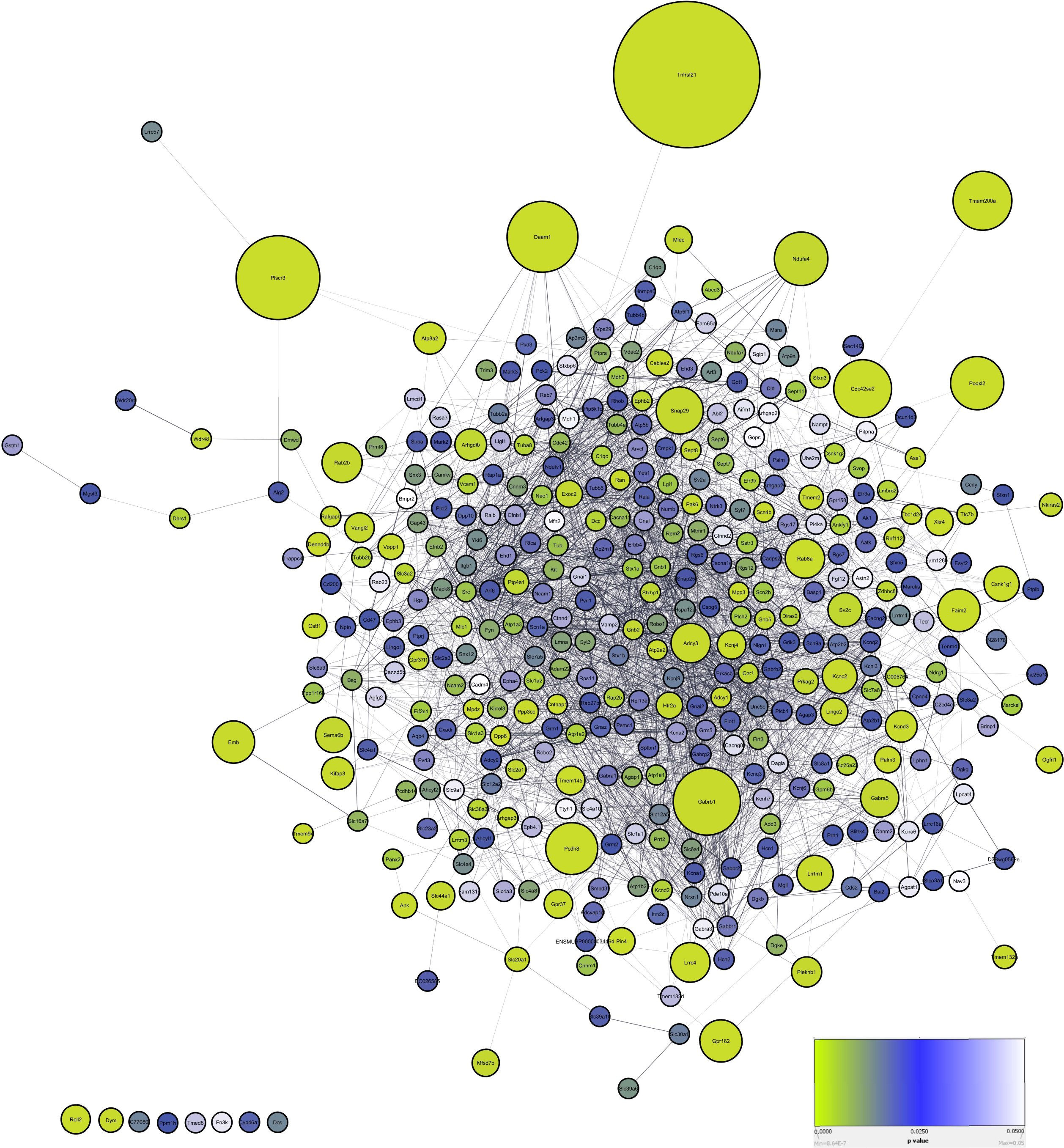
The neuronal ciliary network. This is a clustergram topology of all proteins enriched in SSTR3 iBioID. It is composed of protein networks, with proteins shown as spheres and their interactions as lines. Cytoscape software was deployed to generate this network, relying on its mining capabilities to extensively search the literature. The size of each sphere is correlated to the fold change value and the color shade to the p-value.

One enriched pathway identified in the neuronal ciliary network was axonal guidance signaling **(Figure S3D**, **Table 4)**. Previous studies showed that neuronal cilia signaling is critical for axonal guidance and branching complexity ^24–26,59^. Our results identified 39 proteins involved in axon guidance signaling **(Figure 2C**, **Table 4)**. As part of this module, we isolated the Ephrin/Eph cluster including ephrin ligands, receptors and the roundabout guidance receptors **(Figure 2A**, **4A, and Table 1)**. We assessed the subcellular localization of four proteins, including two ephrin ligands, EFNB1, EFNB2, as well as two Eph receptors, EPHA4, and EPHB2, by immunofluorescence labeling in primary cortical neurons. We co-stained the protein of interest with ARL13B and Pericentrin. All four tested proteins exhibit a similar and intriguing ciliary localization of a two-foci pattern along the neuronal cilium (% of positive cilia for EphA4: 92.57% ± 4.28; Efnb1: 72.96% ± 5.97; Efnb2: 85.36% ± 1.69; Ephb2: 26.94% ± 2.92) **(Figure 4B and C)**. In addition, EphA4 displayed a centrosomal localization often accompanied by one dot along the cilium instead of a two-foci pattern (Centrosomal EphA4: 66.24% ±6.84) **(Figure 4B)**. Since the dissociated cultures used to derive mouse primary cortical neurons can be heterogenous, we used the neuronal marker NeuN to distinguish between neuronal cilia and other cell types. Most neuronal cilia were enriched with EphA4, Efnb1, and Efnb2, with a lesser extent for Ephb2. To further confirm these results, we transiently expressed in IMCD3 cells two Eph/Ephrin proteins, EphA4-GFP and EphB3-V5. We next performed immunofluorescence with antibodies to GFP or V5, respectively, with the ciliary markers AC3 or ARL13B and Pericentrin. Consistent with the endogenous EphA4, EphA4-GFP was observed in distinct foci throughout the cilium. However, we detected more than two foci, which may be due to overexpression or a different regulation in the non-neuronal IMCD3 cells. EphB3-V5, on the other hand, exhibited a more uniform ciliary localization compared to other Eph/Ephrin-tested hits **(Figure 4D)**.

**Figure 4:**
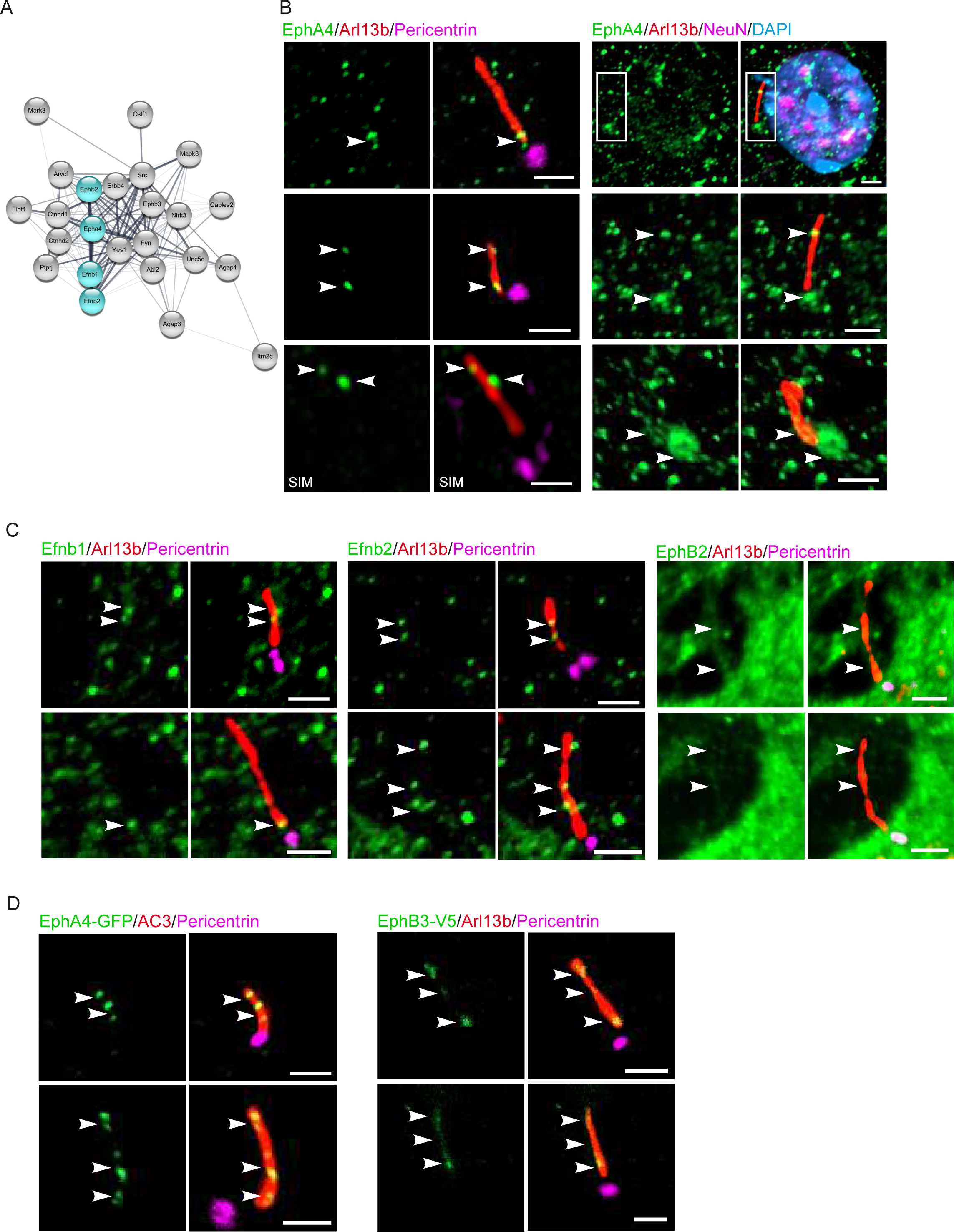
A distinct Eph/Ephrin signaling node is enriched in neuronal cilia. **A.** Clustergram topology of Eph/Ephrin proteins enriched in the cilia iBioID dataset The highlighted hits are validated in the following sections of this figure. **B.** Left panel: Representative immunofluorescence images acquired with the Structured illumination microscopy (SIM). Wild type primary cortical neurons are labeled with EphA4 (green) localizing to cilia co-stained with the ciliary marker Arl13bb (red) and centrosomal marker Pericentrin (magenta). Right panel also shows several examples of EphA4 ciliary enrichment (green) co-labeled with Arl13b and the neuronal marker, NeuN (magenta). DNA is stained with DAPI (blue). Arrows indicate areas of colocalization of EphA4 and Arl13b or Pericentrin. Scale bars: 1 µm. **C.** Representative immunofluorescence images of Efnb1, Efnb2, and EphB2 (green) enriched in neuronal primary cilia labeled with the ciliary marker Arl13b (red) and centrosomal Pericentrin (magenta). Arrows indicate the colocalization of Ephrin/Eph components with Arl13b. Scale bars: 1 µm. **D.** Representative immunofluorescence images of overexpressed EphA4-GFP and EphB3-V5 (green) in IMCD3 cells. Primary cilia are labeled with the ciliary marker AC3 or Arl13b (red) and centrosomal marker Pericentrin (magenta). Arrows show ciliary enrichment of EphA4-GFP and EphB3-V5. Scale bars: 1 µm.

The gamma-aminobutyric acid (GABA) is a major inhibitory neurotransmitter in the mammalian brain ^60^. We identified GABA receptor signaling as the top pathway enriched in our dataset, uncovering 27 components in our neuronal cilia dataset **(Figure 2D**, **S3D and Table 4)**. Our cilia dataset contained 8 sub specific GABA receptors that include the GABA-A receptors, GABRG2 and GABRA1, and the GABA-B receptor, GABBR1. We evaluated the subcellular localization of these proteins by immunofluorescence in mouse primary cortical neurons. GABRG2 showed a consistent, punctate signal throughout the cilium (GABRG2: 77.92% ± 3.52) **(Figure 5A)**. GABRA1: 59.12% ± 3.84) and GABBR1 (48.74% ± 3.13) exhibited a lower number of foci throughout the cilium **(Figure 5A and B)**. We additionally assessed a GABA transporter, SLC6A1, that can mediate GABA levels in extracellular spaces such as the synaptic cleft. We found that this transporter localized to neuronal cilia (SLC6A1: 51.47% ± 2.30) **(Figure 5C)**.

**Figure 5:**
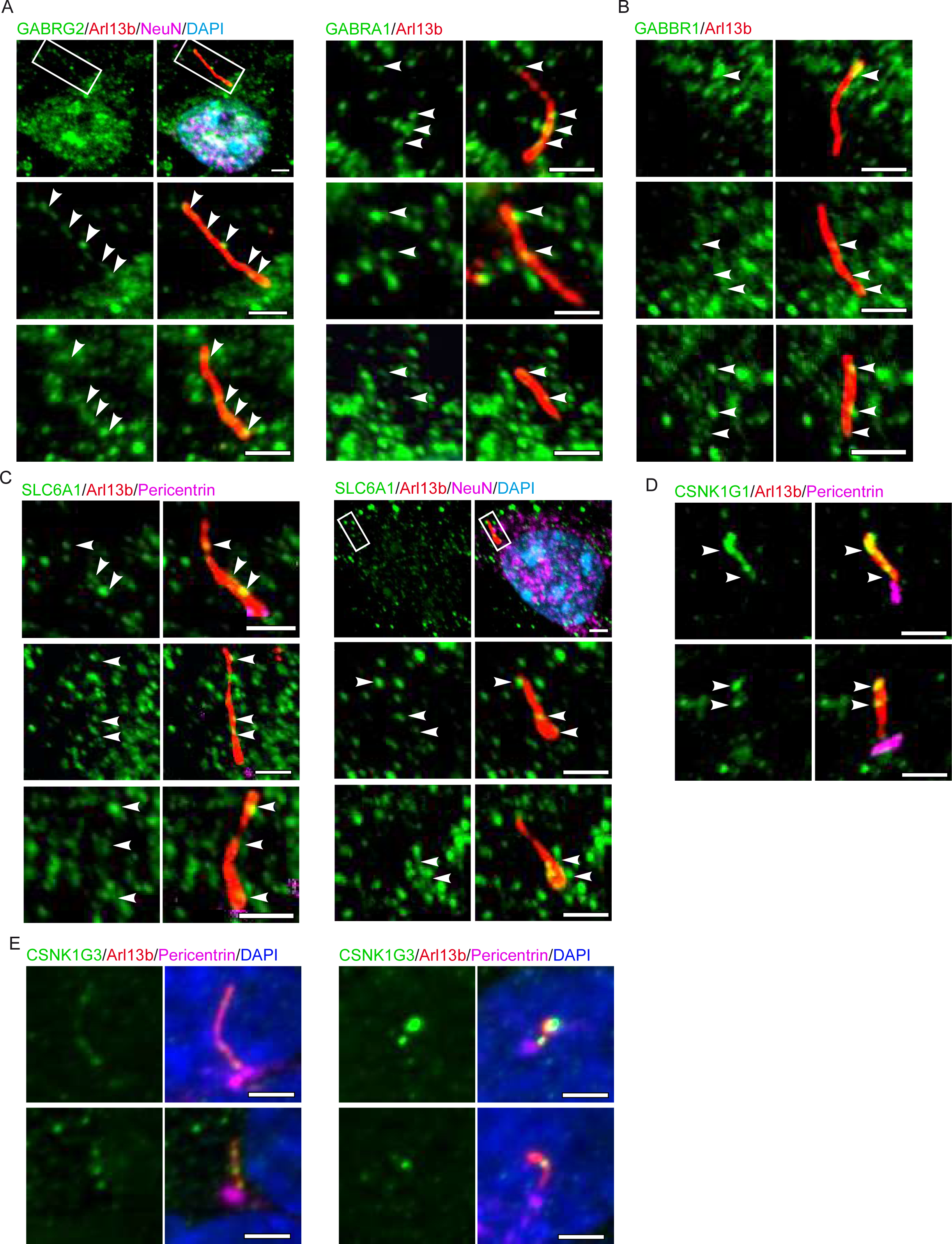
GABAergic and CK1 proteins are enriched in neuronal cilia. **A.** Representative immunofluorescence images of GABRG2 or GABRA1 (green) localizing to neuronal primary cilia labeled with Arl13b (red). Wild type neurons are labeled with and identified by NeuN (magenta). Arrows indicate areas of colocalization of GABRG2 or GABRA1 and Arl13b. Scale bars: 1 µm. **B.** Representative immunofluorescence images of GABBR1 (green) enriched in neuronal primary cilia stained with Arl13b (red). Arrows denote enrichment of GABBR1 in neuronal cilia. Scale bars: 1 µm. **C.** Several examples of representative immunofluorescence of SLC6A1 (green) faintly enriched in neuronal primary cilia. White box denotes the ciliary region shown in the lower section. Arrows are ciliary regions with SLC6A1 enrichment within the neuronal cilium. Scale bars: 1 µm. **D.** CSNK1G1 and **E.** CSNK1G3 are labeled in primary cortical neurons and IMCD3 cells, respectively, with Arl13b (red) and Pericentrin (magenta). DNA was stained with DAPI (blue).

Lastly, we have found that two members of the highly conserved Casein Kinase 1 (CK1) superfamily, CSNK1G1 and CSNK1G3, were significantly enriched in most primary cilia **(Figure 5D and E)**. CK1 is known to modulate fast synaptic transmission, mediated by glutamate, the brain excitatory neurotransmitter ^61^. Overall, the subcellular localization of tested hits showed discrete enrichment throughout the neuronal cilium. This confirms that iBioID successfully identified novel ciliary regulators as a promising first step to defining the ciliary signaling in the brain.

### Cilia iBioID underscores ciliary links of neurological disorders

To uncover new putative linkages between neuronal cilia and human neurological disorders, we again employed the IPA database **(Table 1)**. We found that 199 of the cilia-enriched hits are mutated in or associated specifically with neurological disorders (IPA, p-value range 6.65e-6 – 4.03e-28). In contrast, the background hits depleted in SSTR3 iBioID (or enriched in BioID2 dataset) were not associated with brain disorders but rather cancer (120 molecules, IPA, p-value range 1.8e-2 – 2.37e-28). We provide a comprehensive analysis of the cilia dataset, shown as categories of diseases and functions accompanied by their p-values **(Table 4)**. From these categories, we chose key diseases/functions and overlapped their associated proteins into our neuronal ciliary network to generate clustergrams **(Figure 3 and Figure 6).** About 95 genes were linked to movement disorders and motor dysfunction **(Figure 6A and Table 5)**, which was previously observed in several ciliopathies and cilia-related spinocerebellar ataxia ^11,16,24^. A recent study demonstrated that neuronal primary cilia are disrupted in the dentate gyrus of Fragile X Syndrome, which is the most common monogenic cause of autism spectrum disorder ^62^. We therefore overlapped our dataset with the SFARI Autism database (https://gene.sfari.org/database) of genes directly linked to the autism spectrum disorder **(Figure 6B)**. We found 64 genes present in our neuronal cilia dataset, indicating a deeper yet underexplored linkage between autism and primary cilia function in the brain. Similarly, we uncovered 43 genes that were related to disorders with cognitive impairment **(Figure 6B)**, considered one of the cardinal features of ciliopathies ^22^. Increasing evidence highlights defects in primary cilia structure and function in neurodegenerative disorders. These connections have been reported particularly for Alzheimer’s, Huntington’s, and Parkinson’s disorders ^63,20,21,18,64^, however, the cellular and molecular mechanisms underlying these links have yet to be uncovered. Our findings revealed for the first time that many molecules associated with these neurodegenerative disorders likely reside in the ciliary membrane of neurons **(Figure 6C)**. Altogether, our results constitute a foundation of new information about the molecular composition of neuronal cilia with novel connections to brain disorders.

**Figure 6:**
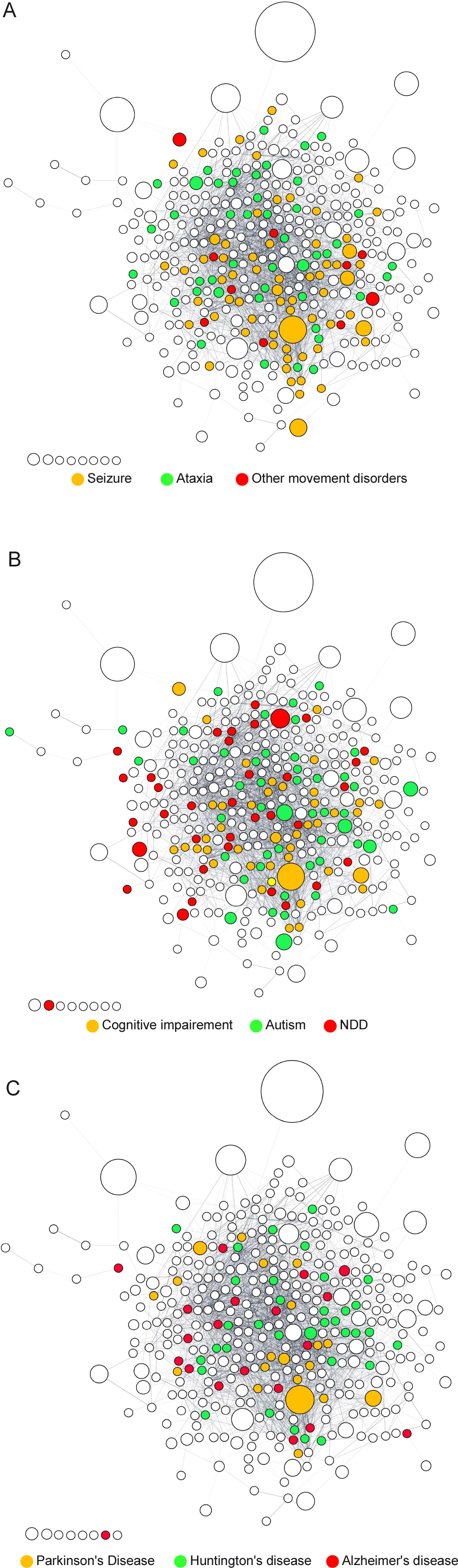
Insights into connections between neuronal cilia and brain disorders. **A.** Clustergram topology of hits associated with cognitive impairment (orange), autism (green), and neurodevelopmental disorders (NDD, red) **B.** Clustergram topology of proteins associated with seizure (orange), ataxia (green), and other movement disorders (red) **C.** Clustergram topology of hits associated with Parkinson’s disease (orange), Huntington’s disease (green), and Alzheimer’s disease (red). For all clustergrams, overlapping hits were shown in one of the categories.

## Discussion

To begin dissecting how cilia mediate signaling events important for neural function, we applied iBioID as a strategy to capture the specific protein repertoire of neuronal cilia in the mouse brain for the first time. Our approach represents a successful adaptation of iBioID to characterize low-abundance cellular structures and organelles, such as the primary cilium, wherein only one is present on each neuron. This contrasts with synaptic compartments, which are present in thousands per neuron. Using a membrane protein as bait, most of our ciliary hits were membrane-related proteins and did not systemically cover other ciliary compartments, including the ciliary barrier and the transition zone. Key ciliary cargoes, such as IFT and BBS complexes, were also not detected. A holistic strategy is therefore needed that combines several baits to achieve full coverage of the protein content of the neuronal cilium. Although cilia iBioID did not comprehensively cover every ciliary sub-compartment, we identified novel neural-specific proteins forming intriguing ciliary enrichment that might be of relevance to brain architecture and function. In most cases, we noticed discrete foci along the cilium, sometimes with similar intensity to non-ciliary localization. Recent discoveries showed that neuronal cilia are part of a tetrapartite synapse composed of pre- and post-synaptic compartments and astrocytes ^30^. We speculate that ciliary Eph/Ephrin and GABAergic signaling components can be among the molecular effectors potentially acting as focal points for cellular interaction, favoring privileged cell-cell communication and signaling. This, in turn, might be key to modulating brain function and behavior.

Our cilia iBioID is a critical first step to understanding the molecular mechanisms by which neuronal cilia mediate neural function. Cilia in the brain are structurally diverse and significantly longer than those of most immortalized cell lines. This is true for different cell types, such as neuronal and glial cells. For example, astrocytic cilia seem to be embedded in pockets, unlike neuronal cilia. They also show diverse interactions with the cortical connectome ^30,31^. These differences suggest functionally distinct roles that call for extensive defining of proteomes of distinct neural and glial cell types. It will also be crucial to define the protein composition of brain cilia in ciliopathies and neurological disorders, which is expected to unveil ciliary perturbations and their effects on neural signaling and circuits.

We found the GABA inhibitory pathway to be our top enriched pathway. Recent studies showed that acute disruption of ARL13B enhances the excitatory capacity of pyramidal neurons ^65^. This may indicate an inhibitory role for primary cilia, which is consistent with our findings. It is likely that specific focal points on neuronal cilia either release or receive GABAergic signaling, which in turn may lead to intracellular changes within the receiving neuron. It is also plausible that glial cells may also be involved in these cell-cell interactions. Recent studies have reported that neuronal cilia can play a role in cell-cell communication between neurons and astrocytes. The absence of cilia in neurons in the brain led to significant changes in astrocytes, which became reactive and showed a reduction in their intracellular lysosomal level ^66^. It is still unclear how astrocytes sense the absence of cilia in neurons. Astrocytes are ciliated but are inclined to become non-ciliated in aged mouse brains ^66^. The exact roles of astrocytic cilia in the brain remain undefined. Altogether, all these findings indicate the complexity of the neural tissue and the likelihood that cilia could function as a novel inter-cellular lever for communication between brain cells. The cilium may also act as a gauge of brain health and age and potentially as an early pathological sign of cell-cell communication disruptions.

The neuronal cilium has been recently linked to serotoninergic signaling and reported to form a novel type of synapse called the axo-ciliary synapse. Brainstem serotonergic axons release serotonin directly into cilia of the hippocampal CA1 pyramidal neurons, which leads to changes in chromatin accessibility ^29^. In our dataset, we only identified one receptor for serotonin, HTR2A. Our cilia iBioID dataset does not have cell-type level resolution to distinguish between neuronal sub-types and their associated neurotransmitters, including serotonin, dopamine, and glutamine. Further studies targeting and comparing cilia of distinct populations of neurons are key to building a comprehensive foundation of primary cilia biology in the brain. Our finding is a proof-of-principle that further screening of the neuronal ciliome will be feasible in the brain. Our cilia iBioID will allow future studies to identify the molecular dysfunctions within neuronal cilia found in ciliopathies and neurodegenerative disorders. These future directions will bring greater understanding about cilia pathogenicity and how it could affect neural function and, in turn, lead to neurological deficits.

This research lays the framework for understanding the intrinsic signaling transmitted by the neuronal cilium, as well as potential external stimuli and downstream signals that influence brain function and homeostasis. The long-term goal would be to develop unique foundations for cilia biology in the brain by connecting novel neural ciliary signaling to subtypes of neurons and their specialized circuits. Ultimately, using the neuronal cilium as a targetable molecular lever in a novel therapeutic strategy to help correct neurological deficits in human disease.

## Supplementary figures

**Figure S1:**
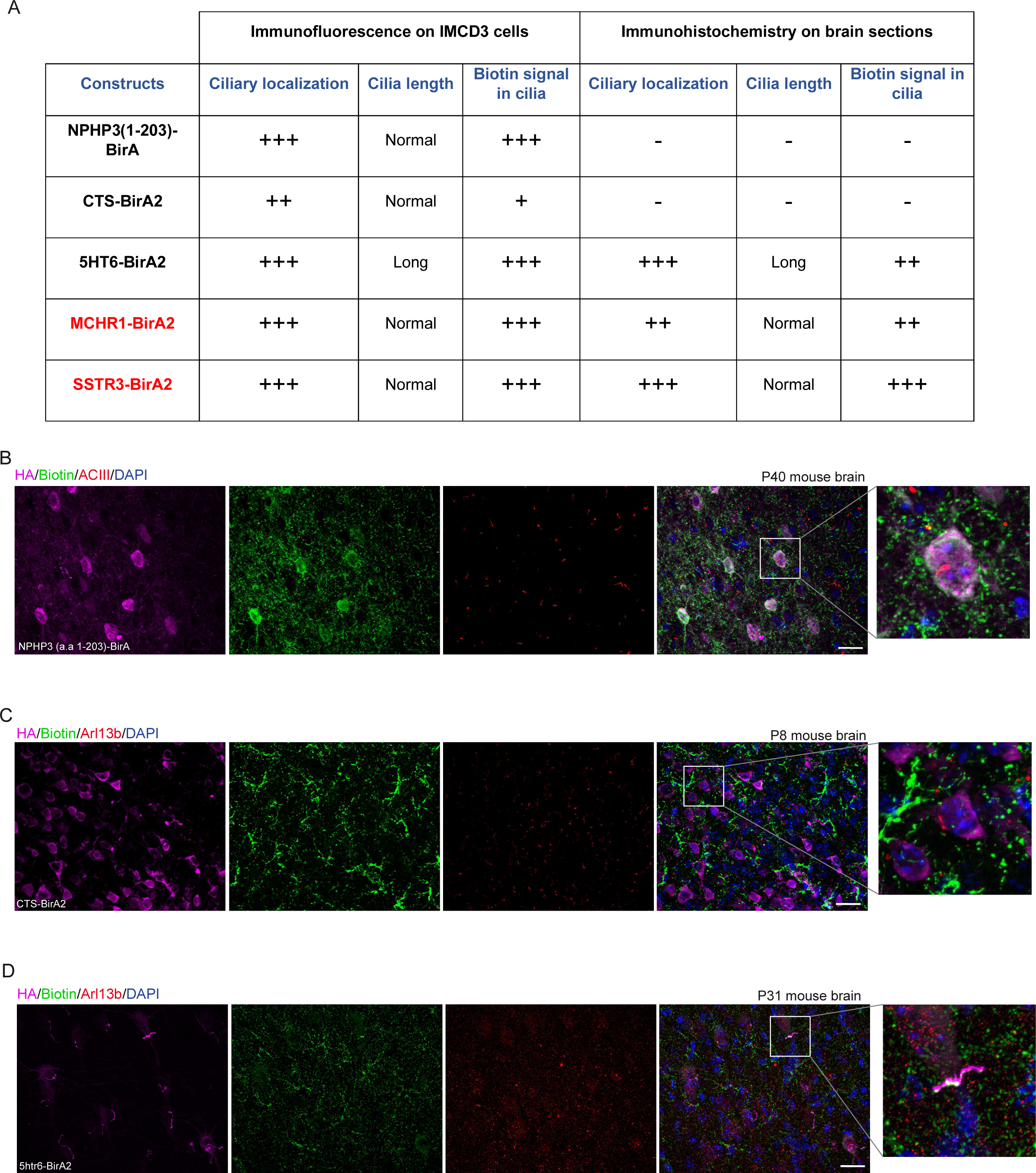
Testing of ciliary baits to target biotin ligase. **A.** A summary of tested ciliary baits to target the biotin ligase to neuronal cilia in vivo. Baits were first tested in IMCD3 cells and then assessed in the mouse brain, as detailed in figure 1A. Immunohistochemical labeling from mouse brain sections transduced with **B.** AAV-NPHP3 (a.a 1-203)-BirA. **C.** AAV-ciliary targeting sequence of Fibrocystin (CTS)-BirA2. **D.** AAV-5HTR6-BirA2. Sections are labeled with antibodies to Biotin (green), ciliary markers ACIII and Arl13b (red), and HA (magenta). DNA was stained with DAPI. Scale bars: 20 µm.

**Figure S2:**
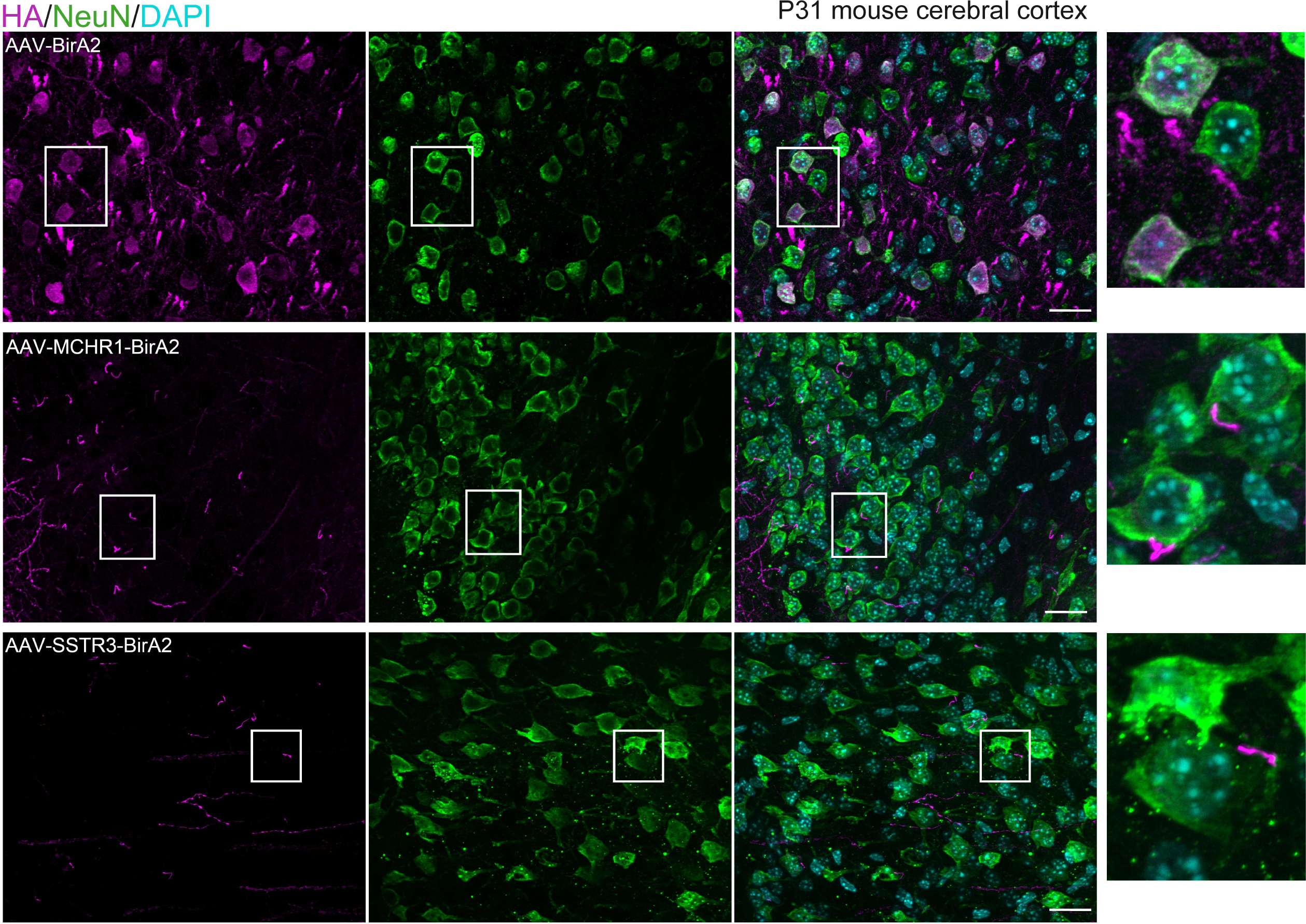
BirA2 constructs are exclusively expressed in neurons. Immunohistochemical labeling from mouse cerebral sections transduced with AAV-BirA2 or AAV-GPCRs-BirA2. Sections are labeled with antibodies to HA (magenta) and the neuronal marker NeuN (green). DNA was stained with DAPI. Scale bars: 20 µm.

**Figure S3:**
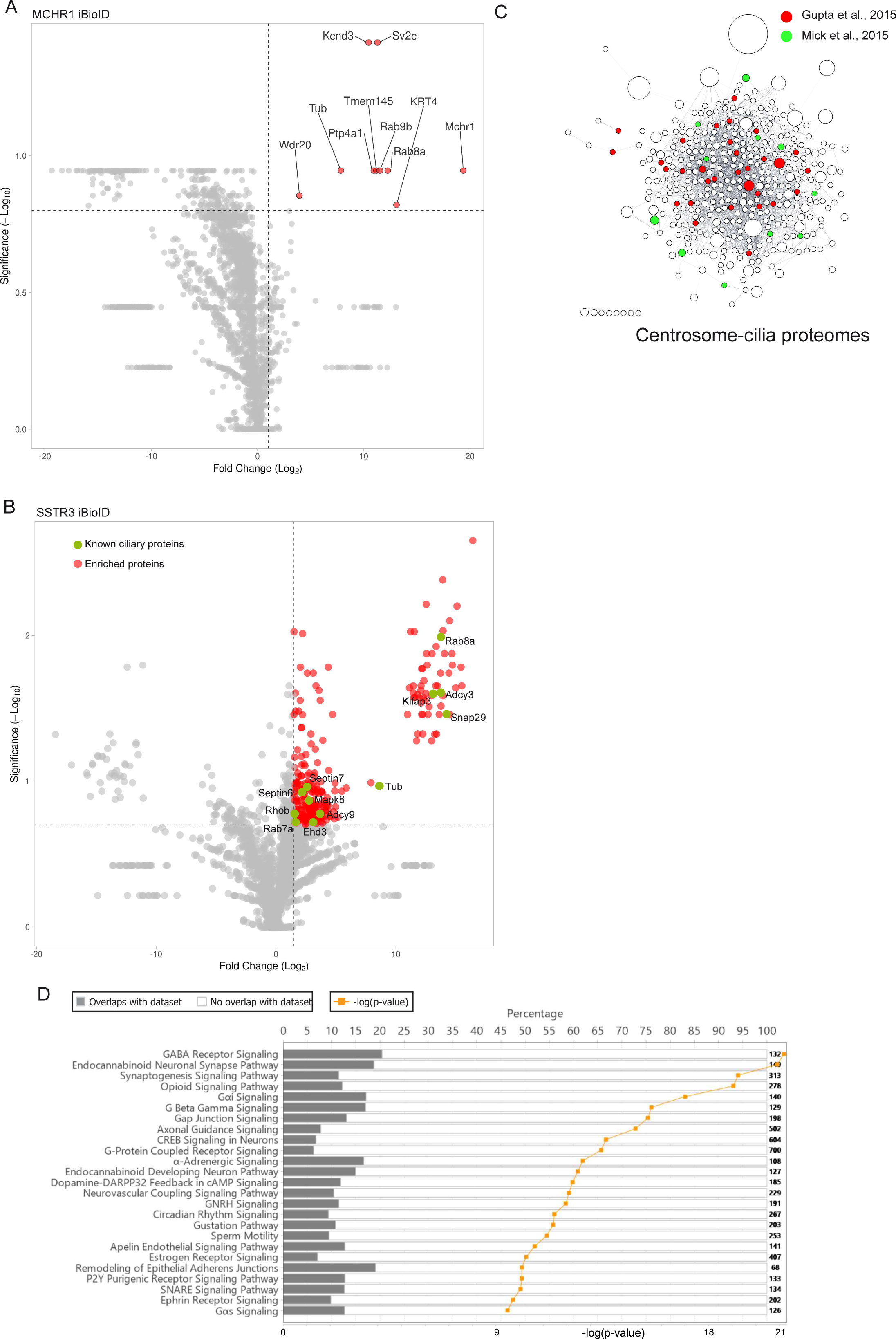
Volcano plots of SSTR3 and MCHR1 BioID and GO analysis. **A.** Volcano plot of MCHR1-proximate proteins labeled by BioID. The red dots show the proteins considered hits, delimited by the selected thresholds: log2 fold change 2 and significance 0.7. The X-axis denotes the log2 fold change of MCHR1-BirA2: BirA2 control. In the Y-axis, significance displays the negative log10 transformed p-value for each protein. **B.** Volcano plot of SSTR3-proximate proteins labeled by BioID as shown in figure 2A. The green dots highlight known ciliary proteins overlapping with cilia proteomes discovered in Gupta et al., 2015 and Mick et al., 2015. **C.** Clustergram topology of centrosome-cilia proteomes (Gupta et al., 2015 (red) and Mick et al., 2015 (green)) enriched in SSTR3 iBioID dataset. **D.** Gene ontology analysis of hits enriched in SSTR3 iBioID. The table summarizes the most enriched signaling pathways with their corresponding p-values.

---

**Table 1:** Quantitative proteomics results of control and GPCRs iBioID.

**Table 2:** Gene ontology analysis of SSTR3-iBioID hits showing categories of protein functions and their corresponding p-values within the dataset.

**Table 3:** Analysis of Diseases and functions categories of hits enriched in SSTR3-iBioID dataset.

**Table 4:** Most enriched canonical pathways that overlap with our SSTR3-iBioID.

**Table 5:** An exhaustive list of disease categories and their associated genes, enriched in the SSTR3-iBioID dataset.

## Acknowledgments

We thank Dr. Shataakshi Dube O’Neil and Dr. Erin Spence for their technical advice and assistance. We also thank Dr. Erik Soderblom and the Duke Proteomics and Metabolomics Core for their help with the cilia iBioID. This work was supported by a Duke University Chancellor’s Discovery Award to SCG, and by National Institutes of Health awards R00 HD076444 and R01 HD099784 to SCG; and R35GM151229 (NIGMS, NIH) to AL. The Regeneration Next Initiative at Duke University partially funded AL research with the RNI Postdoctoral Fellowship.

## Materials and methods

### Animals

FVB/NJ mice (stock no. 001800) were obtained from Jackson Laboratory. Rosa26-LSL-Cas9 knockin (stock No. 026175) mice were obtained from Jackson Laboratories. Transgenic mice expressing Arl13b-mCherry were a gift from Dr. Kathryn Anderson. Mice were housed (up to 5 mice per cage) in the Division of Laboratory Animal Resources facility at Duke University. All the experimental procedures have been approved by the Duke Institutional Animal Care and Use Committee (IACUC) and followed the National Institutes of Health guidelines.

### Cloning and IMCD3 culture

cDNAs for mouse GPCRs, SSTR3, or MCHR1 cDNAs were cloned into the pAAV-BirA2-HA-linker-BirA2-HA (gift from Dr. Scott Soderling, Duke University). BirA2-HA was C-terminally fused to either SSTR3 or MCHR1. Similarly, BirA was fused to NPHP3 (a.a 1-203) fragment. Plasmids containing gene sequences for EphA4 (pDONR223-EPHA4, Addgene, 23919) and EphB3 (FUW-ubiquitin-EphB3-SV40-GFP, Addgene, 65443), two ciliary proteome candidates, were obtained and used to perform gateway recombination cloning. pDONR223-EPHA4 was directly cloned into PFLAP-DEST using the LR Clonase reaction (ThermoFisher, 11791020). FUW-ubiquitin-EphB3-SV40-GFP underwent PCR amplification and purification using the following primer sequences to amplify the EphB3 sequence: forward 5’CACCATGGCCAGAGCCCGCC 3’, reverse 5’ TCAGACCTGCACAGGCAGCGT 3’. With the PCR purified product, TOPO cloning was performed (ThermoFisher, K240020). Then, pENTR/D-TOPO-EphB3 was cloned into pcDNA3.1/nV5-DEST (ThermoFisher, 12290010). After amplification of the final cloned constructs, transient transfection was performed in mouse inner medullary collecting duct cells (mIMCD-3) (ATCC, CRL-2123). These were plated on 15 mm coverslips and maintained in DMEM: F-12, HEPES (ThermoFisher, 11330057) supplemented with 10% v/v heat-inactivated fetal bovine serum (ThermoFisher, 10082147). Transient transfection was performed accordingly with Lipofectamine 2000 Transfection Reagent (ThermoFisher, 11668019).

### AAV production and purification

For each AAV virus, we transfected six 15 cm plates of HEK293T cells with pAd-DeltaF6, pAAV serotype 2/9, and pAAV-GPCR-BirA2-HA or pAAV-BirA2-HA. HEK293T cells were cultured in DMEM with 10% fetal bovine serum. We used 7,5 mM polyethylenimine (PEI) and incubated transfection complexes for 18 hours. Growth media was replaced the next day. After 24 hours, cells and supernatants are collected and centrifuged at room temperature for 5 min at 12000 rpm. Pellets were then resuspended in 4 ml of cell lysis buffer (15 mM NaCl, 5 mM Tris-HCl, pH 8.5). We performed three cycles of freeze-thawing in a dry ice-ethanol bath and a 37°C water bath. Benzonase (50U/ml) was added to cell lysates and incubated for 30 minutes in a 37°C water bath. Lysates were centrifuged at 4500 rpm for 30 minutes at 4°C. Supernatants were collected and added on a gradient of 15%, 25%, 40%, and 60% of iodixanol solution. We centrifuged at 30,000 rpm for 2 hours to carefully collect the viral solution. We concentrated AAV viruses using Amicon filters and washed them four times with 1X PBS. We finally resuspended them in 100 l. To inject equal amounts of viruses, we tittered viral solutions by quantitative PCR. Until needed, AAVs were stored at - 80°C.

### *In vivo* BioID of neuronal cilia

#### AAV intracranial injections in P0-P2 pups

AAV viruses coding for BirA2, MCHR1-BirA2 or SSTR3-BirA2 were produced, purified, and titrated. Intracranial injections of AAVs were performed in P0-P2 brain pups. Note that wild-type FVB/NJ background was used for the quantitative proteomics and AC mice for immunohistochemical labeling. For optimal labeling throughout the forebrain, we injected ice-anesthetized pups with 1.5 l - 2.5 l per hemisphere of concentrated AAVs diluted in 1X PBS (from 10^10^ to 1.5 10^11^ virus/ l) using a 10 l Hamilton syringe and a 1-inch needle (33 gauge needle with a 20 degree bevel). A heating pad was used to facilitate the recovery of injected pups until their movement pattern and skin color were back to normal. After injections, the whole litter was reintroduced to the dam and closely monitored for at least 24 hours.

### Biotin labeling in mice

At P19-P25, we subcutaneously injected 1 ml of 6.66 M of biotin diluted in 1X PBS (sterile) once a day for 5 days. We continued subcutaneous injections that we combined with the intraperitoneal injections (1 ml per day) for 6 days. The two injections were performed at different times during the day for the welfare of the animal. AAV-BirA2-, MCHR1-BirA2, and SSTR3-BirA2-injected brains were carefully extracted at P31-P36 of age. The cerebellum that showed weak biotin labeling was removed. The rest of the brain was snap-frozen in liquid nitrogen until pulldowns were performed.

#### Pulldowns of biotinylated protein from brains

##### A critical note

Throughout the purification step, personal protective equipment, including a mask, a clean coat, gloves, and a bouffant cap, should be worn to diminish human keratin contamination. All solutions were also freshly prepared in similar conditions. The quantitative proteomics experiment was performed with AAV injections of ten different litters on different days. For each biological replicate, we used 7 brains per condition that we pooled and crushed into fine powder with a mortar and pestle dipped in liquid nitrogen. The lysis buffer was freshly prepared. Its composition is as follows: [1% NP-40 (85124, ThermoFisher), 2 protease inhibitor tablets (Roche), 20 M MG132 (Sigma-Aldrich), 50 mM Beta-glycerol phosphate (Sigma-Aldrich), and 1:1000 benzonase nuclease (250 U/ L) from Sigma-Aldrich]. Brain powder was put into 10 ml of lysis buffer in Corning 15-ml tubes on ice for each condition. We incubated the extracts on ice for 30 minutes to help lyse the brain powder, followed by a sonication step: 8 times for 3 seconds with an amplitude of 25%. We then added 1 ml of 10% SDS and incubated for 15 minutes on ice. Extracts were transferred to 1.5 ml low-retention tubes or ultracentrifuge tubes and incubated for an additional 15 min on ice. Lysates were then centrifuged at 15000 rpm for 45 min at 4°C and the supernatants were transferred to new 15-ml tubes (Corning). We next washed the Neutravidin beads five times with lysis buffer with additives. We added 120 l of washed Neutravidin beads to each supernatant and incubated each pulldown at 4 °C with tube rotation at a speed of 25 rpm. The next day, we centrifuged samples at 3000 rpm for 3 min to recover the beads and remove the supernatant. The final step consisted of several washes sequentially performed on beads. We carried out 3 washes with 12 ml of each of the following solutions: (1) 2% SDS; (2) 1% Triton, 1% deoxycholate, and 25 mM LiCl; (3) 1 M NaCl ^39^. For each wash, we rotated samples for 5 minutes at room temperature with a 30 rpm rotation speed. We then spun at 3000 rpm for 1 min after each wash. Next, we washed 4 times with 50 mM ammonium bicarbonate. Beads were transferred to low-retention 1.5 ml tubes. 220 l of 2X Laemmli supplemented with 5 mM biotin were added, and samples were incubated for 8.5 min at 95 ° C. To remove any residual beads, we centrifuged twice and recovered about 160 l from each pulldown. We kept 20 l to validate the level of biotin labeling for each sample by western blot. Samples were stored at - 80° C until submitted for sequencing with mass spectrometry. We carried out western blots to assess the overall biotin labeling and a ciliary marker, ARL13B.

### Quantitative proteomics

#### Sample Preparation

The Duke Proteomics Core Facility (DPCF) received 9 samples (3 replicates of BirA2, 3 of MCHR1, and 3 of SSTR3). Samples were supplemented with 160 l of 10% SDS in 50 mM TEAB, then reduced with 10 mM dithiolthreitol for 30 min at 80C and alkylated with 20 mM iodoacetamide for 30 min at room temperature. Next, they were supplemented with a final concentration of 1.2% phosphoric acid and 2,730 L of S-Trap (Protifi) binding buffer (90% MeOH/100mM TEAB). Proteins were trapped on the S-Trap, digested using 20 ng/ l sequencing grade trypsin (Promega) for 1 hr at 47C, and eluted using 50 mM TEAB, followed by 0.2% FA, and lastly using 50% ACN/0.2% FA. All samples were then lyophilized to dryness and resuspended in 12 L 1%TFA/2% acetonitrile containing 12.5 fmol/ L yeast alcohol dehydrogenase (ADH_YEAST). From each sample, 3 L was removed to create a QC Pool sample which was run periodically throughout the acquisition period.

#### Quantitative Analysis, Methods

Quantitative LC/MS/MS was performed on 4 L of each sample, using a nanoAcquity UPLC system (Waters Corp) coupled to a Thermo Fusion Lumos high resolution accurate mass tandem mass spectrometer (Thermo) via a nanoelectrospray ionization source. Briefly, the sample was first trapped on a Symmetry C18 20 mm × 180 m trapping column (5 l/min at 99.9/0.1 v/v water/acetonitrile), after which the analytical separation was performed using a 1.8 m Acquity HSS T3 C18 75 m × 250 mm column (Waters Corp.) with a 90-min linear gradient of 5 to 30% acetonitrile with 0.1% formic acid at a flow rate of 400 nanoliters/minute (nL/min) with a column temperature of 55C. Data collection on the QExactive HF-X mass spectrometer was performed in a data-dependent acquisition (DDA) mode of acquisition with a r=120,000 (@ m/z 200) full MS scan from m/z 375 – 1500 with a target AGC value of 2e5 ions followed by 30 MS/MS scans at r=15,000 (@ m/z 200) at a target AGC value of 5e4 ions and 45 ms. A 20s dynamic exclusion was employed to increase depth of coverage. The total analysis cycle time for each sample injection was approximately 2 hours.

Following 13 total UPLC-MS/MS analyses (excluding conditioning runs, but including 4 replicate QC injections; **Table 1**), data was imported into Proteome Discoverer 2.2 (Thermo Scientific Inc.), and analyses were aligned based on the accurate mass and retention time of detected ions (“features”) using Minora Feature Detector algorithm in Proteome Discoverer. Relative peptide abundance was calculated based on area-under-the-curve (AUC) of the selected ion chromatograms of the aligned features across all runs. The MS/MS data was searched against the SwissProt *Mus musculus* database (downloaded in April 2018) with additional proteins, including yeast ADH1, bovine serum albumin, as well as an equal number of reversed-sequence “decoys”) false discovery rate determination. Mascot Distiller and Mascot Server (v 2.5, Matrix Sciences) were utilized to produce fragment ion spectra and to perform the database searches. Database search parameters included fixed modification on Cys (carbamidomethyl) and variable modifications on Meth (oxidation) and Asn and Gln (deamidation). Peptide Validator and Protein FDR Validator nodes in Proteome Discoverer were used to annotate the data at a maximum 1% protein false discovery rate.

### GO and cluster analysis

Gene ontology analysis was performed using Ingenuity Pathway Analysis. We followed the manufacturer’s guidelines to extract the enriched molecular functions, enriched signaling pathways, and associations with diseases. Cytoscape 3 was employed to reconstitute the molecular networks of our cilia iBioID hits. We used the string app, in which we selected a full-string network with a confidence score cutoff of 0.3. Once the network is formed, we display the fold change as the size of the spheres and the p-values with a colored continuous mapping. To highlight certain proteins, we uploaded a data table file as node table columns that we matched with the display name (case-insensitive). We then selected these hits and filled the spheres with the appropriate colors.

### Western blot analysis

Lysates were incubated at 95°C for 5 min. Denatured proteins were separated by SDS-PAGE and transferred onto a PVDF membrane. The membrane was saturated in 5% milk PBST (5% dry powdered milk, 0.1% Tween-20, 1X PBS) for 1 hour and then incubated with primary antibodies overnight. Secondary antibodies (peroxidase-conjugated affinipure or IRDye secondary antibodies). Membranes were then developed with an ECL reagent kit (BioRad) or Li-COR imaging system. The western blots were quantified using Fiji software.

### Primary neuronal culture

Primary neuronal cultures were derived from Cas9 knockin P0 pups. All brain dissections were performed in cold 1 X PBS. After brain extraction, we carefully removed and discarded the meninges. Cortices and/or hippocampi were isolated and cut into pieces before being transferred to 15 cm tubes in 1X PBS. After discarding the PBS, we added a pre-heated dissociation solution (Mix A [40 ml 1X PBS, 10 mg DL-cysteine HCl (Sigma C9768), 10 mg BSA, 250 mg D-glucose (Sigma G6152)]; 5 U/ml Papain; and 120 U/ml DNase). Dissociate for 30-40 minutes at 37 °C. We then replaced the solution with neuronal media (50 ml neurobasal, 1 ml neuronal B27 supplement, 500 l Glutamax, and 60 l Pen/Strep). Mechanical dissociation is carefully performed using a 1 ml micropipette (10 times). Cells were plated on pre-coated poly-D-lysine (0.1 mg/ml, 37° C; 5% CO2 for at least 2 hours) coverslips in a 24-well plate. Cells were fixed at DIV10-15 with 4% paraformaldehyde for 15 min and permeabilized with cold 100% methanol for 5 min.

### Immunofluorescence and microscopy

Neurons were fixed between DIV7 and DIV14 in 4% Paraformaldehyde (PFA)/phosphate-buffered saline (PBS) (Santa Cruz, sc-281692) for 15 minutes at room temperature. Then, they were permeabilized with ice cold methanol (VWR, BDH1135) for 5 minutes at -20 . Neurons were blocked with blocking buffer PBS + 1% BSA and 0.2% Triton X-100 for 1 h at room temperature. Cells were fixed with 4%PFA for five minutes at room temperature and permeabilized with ice cold methanol for 5 minutes at -20 . Cells were blocked with blocking buffer PBS + 1% BSA, 0.2% Triton X-100, and 5% goat serum for 30 minutes at room temperature. Samples were then stained with their respective primary antibodies listed previously overnight at 4 . The next morning, after washing with PBS 3 x 5 min, samples were incubated with appropriate secondary antibodies for 1 h (neurons) or 30 minutes (transfected cells) at room temperature. Samples were then incubated with DAPI (Sigma, D9542, 1:1000) and then washed with PBS 3 x 5 min and mounted with ProLong Gold Antifade Mountant (ThermoFisher, P36930). Mounted coverslips were then imaged on a Zeiss Axio Observer-Z1 widefield microscope equipped with a Zeiss Plan-Apochromat 63x / 1.4 Oil objective, an Axiocam 506 monochrome camera, and an Apotome setting. To accurately quantify the endogenous presence and overexpression of ciliary proteome candidates and ciliary markers, a Z-stack was acquired, a maximum intensity projection for each image was created, and the maximum intensity projections were analyzed using Fiji software.

### Immunohistochemistry

Transcardiac perfusion of P31-P36 mice was performed using 1X PBS followed by 4% paraformaldehyde (PFA). After extraction, the brain was fixed in 4% PFA for 2 hours at 4° C. PFA was washed out with 1X PBS and replaced with 30% sucrose for overnight incubation. The brains were frozen in disposable embedding molds in OCT compound. We then generated coronal and sagittal brain sections of about 20 microns, mounted on charged slides. To permeabilize the tissue, we incubate it with 1% Triton diluted in 1X PBS for 1 hour. The tissue is blocked in blocking buffer (0.1% Triton X-100, 5% FBS (Goat), 1% bovine serum albumin, 0.02% Sodium Azide, 2% FBS, 0.1% Triton X-100 in 1X PBS) for 1 hour. Primary antibodies were incubated for 3 hours at room temperature or overnight at 4 °C, followed by 2 hours of incubation with secondary antibodies (1:500; Alexa Fluor 488, Alexa Fluor 568, Alexa Fluor 647; Molecular Probes, Thermo Fisher Scientific). DNA was stained with DAPI for 1 min. Coverslips were added to glass slides using ProLong Gold mounting media (Molecular Probes, Thermo Fisher Scientific). The stained tissue was imaged with a Zeiss Axio Observer-Z1 widefield microscope equipped with a Zeiss Plan-Apochromat 63x/1.4 Oil objective, an Axiocam 506 monochrome camera, and an Apotome setting. We acquired a Z-stack and used Fiji software to make a maximum intensity projection for each image.

### Primary Antibodies

We used the following antibodies: Anti-ARL13B (IF:1/500, western blot (WB): 1/1000; NeuroMab, 11000053); Anti-HA (clone 3F10, IF:1/1000, 11867423001, Sigma); Anti-NeuN (IF: 1:300, ab177487, Abcam); Anti-V5 (IF: 1:500, R960-25, Invitrogen); Anti-DR6 (TNFRSF21) (clone 6B6, IF: 1:200, MABC1594-25UL, Millipore); Anti-ARL13B (Immunofluorescence (IF): 1:500, Abcam, ab136648); Anti-ARL13B (IF: 1:500, Proteintech, 17711-11AP); Anti-Pericentrin (IF: 1:500, BD Biosciences, 611814); Anti-Pericentrin (IF: 1:500, Abcam, ab4448); Anti-NeuN (IF: 1:500, Abcam, ab104224); Anti-NeuN (IF: 1:500, Abcam, ab177487); Anti-GFP (IF: 1:1000, Invitrogen, A11122); Anti-V5 Tag (IF: 1:500, Invitrogen, R960-25); Anti-Adenylate Cyclase 3 (IF: 1:500, Invitrogen, PA5-35382); Anti-Ephrin-B1 (IF: 1:50, Bio-Techne, AF473); Anti-Ephrin-B2 (IF: 1:50, Bio-Techne, AF496); Anti-EphB2 (IF: 1:50, Bio-Techne, AF467); Anti-EphA4 (IF: 1:50, Proteintech, 21875-1-AP); Anti-CSNK1G1 (IF: 1:50, OriGene, TA806333S); Anti-ROBO2 (IF: 1:100, GeneTex, GTX134119); Anti-UNC5C (IF: 1:100, LSBio, LS-B8305-50); Anti-SEMA6B (IF: 1:100, biorbyt, orb422749); Anti-SLC6A1 (IF: 1:500, Invitrogen, PA5-85766); Anti-GABRG2 (IF: 1:100, Proteintech, 14104-1-AP); Anti-GABBR1 (IF: 1:500, Invitrogen, PA5-27725); Anti-GABRA1 (IF: 1:100, Proteintech, 12410-1-AP), Anti-DR6 (Tnfrsf21) (IF: 1:100, Sigma, MABC1594).

### Secondary Antibodies

We use the following secondary antibodies, and their conditions of use include: Alexa Fluor 488 Goat anti-Rabbit IgG (Invitrogen, A-11008, 1:500); Alexa Fluor 647 Goat anti-Mouse IgG1 (Invitrogen, A-21240, 1:500); Alexa Fluor 647 Goat anti-Mouse IgG2b (Invitrogen, A-21242, 1:500); Alexa Fluor 568 Goat anti-Mouse IgG2a (Invitrogen, A-21134, 1:500); Alexa Fluor 647 Goat anti-Rabbit IgG (Invitrogen, A27040, 1:500); Alexa Fluor 647 Donkey anti-Rabbit IgG (Invitrogen, A-31573, 1:500); Alexa Fluor 568 Donkey anti-Rabbit IgG (Invitrogen, A10042, 1:500); Alexa Fluor 647 Donkey anti-Mouse IgG (Invitrogen, A-31571, 1:500); Alexa Fluor 568 Donkey anti-Mouse IgG (Invitrogen, A10037, 1:500); Alexa Fluor 488 Donkey anti-Goat IgG (Invitrogen, A-11055, 1:500), Alexa Fluor 488 Goat anti-Mouse IgG2a (Invitrogen, A-21131, 1:500), Alexa Fluor 488 Goat anti-Rat IgG (Invitrogen, A-11006, 1:500); Streptavidin, Alexa Fluo 488 (IF: 1:500, S11223, Invitrogen); IRDye 680RD Streptavidin (WB: 1:3000, 926-68079, LiCor).

### Lattice SIM

Primary neuronal cells were plated on high-precision type 1.5 coverslips. We incubated primary antibodies for 24 h at 4°C. We used Alexa secondary antibodies for each staining (IF: 1:500; Alexa Fluor 488, Alexa Fluor 568, Alexa Fluor 647; Molecular Probes, Thermo Fisher Scientific). These were incubated for 1 h 30 min at room temperature. We then mounted the coverslips on glass slides using ProLong Gold mounting media (Molecular Probes, Thermo Fisher Scientific). Our SIM acquisitions were done on a Zeiss Elyra 7 system equipped with lattice SIM technology. The signal was collected using a Zeiss Plan-Apochromat 63/1.4 Oil objective. We kept the laser power and exposure time to a minimum to avoid photobleaching. In the ZEN imaging software, the SIM mode was used to process the acquired images.

### Quantification and statistical analysis

Measurements of cilia rate and candidate endogenous localization frequency or overexpression were performed on at least five randomly selected fields of cells per coverslip for each condition. At least two coverslips were analyzed per condition with similar confluence. The maximum intensity projections were used to perform this analysis with Fiji Software. Analysis was performed with the same signal parameters between candidates and their relative controls.

Endogenous candidate localization frequencies measured on neuronal cilia were determined by concurrent use of a neuronal marker or by morphological recognition of neurons and their cilia. Frequency measurements are an estimation. Localization of candidates was determined by the colocalization of fluorescent signals for the cilia channel and candidate channel. Data are reported as arithmetic means ± SEM, and statistical analysis was performed using paired t-tests and One-way ANOVA with p 0.05 as the cutoff for statistical significance. Statistical analyses were performed using GraphPad Prism 9 software.

